# Bird functional dispersion predicts dengue vector presence across tropical working landscapes in Costa Rica

**DOI:** 10.64898/2026.08.26.747376

**Authors:** Dallas R. Levey, Caroline K. Glidden, Talya Shragai, Andrew J. Chamberlin, Rachel L. Fay, Luis Enrique Chaves-Gonzalez, Jeisson Figueroa-Sandi, Joshua E. Lazaro, Váleri N. Vásquez, Rolando D. Moreira-Soto, Kota Osawa-Pivovarov, Gretchen C. Daily, Diana Rojas-Araya, Adriana Troyo, Erin A. Mordecai

**Affiliations:** Stanford University, Department of Biology, Stanford, California, USA; Stanford University, Natural Capital Alliance, Stanford, California, USA; Stanford University, Doerr School of Sustainability, Stanford, California, USA; Stanford University, Department of Oceans, Stanford, California, USA; Laboratorio de Investigación en Vectores, Centro de Investigación en Enfermedades Tropicales, Universidad de Costa Rica, San José, Costa Rica; Sección de Entomología Médica, Facultad de Microbiología, Universidad de Costa Rica, San José, Costa Rica; Organization for Tropical Studies, Estación Biológica Las Cruces, San Vito, Costa Rica; Stanford University, Department of Biomedical Data Science, Palo Alto, California, USA

**Keywords:** *Aedes aegypti*, *Aedes albopictus*, bioindicators, Costa Rica, dengue, disease ecology, functional dispersion

## Abstract

Identifying reliable indicators of dengue vector habitat is critical for proactive surveillance and spatially targeted vector control. Bird communities are sensitive to habitat quality and landscape condition, and their functional diversity may track ecological conditions that support Aedes mosquito populations. We tested whether bird functional diversity predicts dengue vector presence across two working landscapes in Costa Rica, comparing bird-based predictors against landscape composition, vegetation structure, and plant diversity. Bird functional dispersion (FDis), the spread of species in multidimensional trait space, was the top-ranked predictor for *Ae. aegypti* and *Ae. albopictus* larval presence and for adult *Ae. aegypti*, outperforming all other predictor classes. Probability of presence declined with increasing FDis, where functionally homogeneous bird communities, characteristic of disturbed landscapes, co-occur with higher vector presence, consistent with shared habitat filtering. Functional diversity estimations may offer a practical signal for identifying habitats at elevated dengue vector risk.

## Introduction

Dengue is one of the world’s fastest-growing public health threats, infecting an estimated 400 million people annually and expanding rapidly across the tropics and subtropics (Bhatt et al. 2013; Childs et al. 2025; Hidalgo et al. 2026; Stanaway et al. 2016). While dengue is canonically characterized as an urban disease, rural transmission is increasingly recognized across the Americas and Asia (Man et al. 2023; Shepard et al. 2016; Skinner et al. 2023), where heterogeneous land use and co-occurring vector species complicate risk assessment and vector control. In tropical working landscapes, the ecological drivers of the distributions of the two primary dengue vectors, *Aedes aegypti* and *Aedes albopictus,* are poorly understood, in part because these species respond to fine-scale habitat conditions and availability of anthropogenic water-holding containers that are difficult to characterize from land cover maps alone (Glidden et al. 2026). Improving the ecological predictability of vector presence is a prerequisite for proactive, spatially targeted surveillance and intervention, particularly outside of urban areas.

Biodiversity has been tied to disease risk (Glidden et al. 2021; Keesing et al. 2010; Keesing & Ostfeld 2021; Merrill et al. 2025), though its generality across disease systems remains debated (Salkeld et al. 2013). These associations may reflect direct, causal effects of biodiversity on transmission dynamics, or shared (i.e., correlated) responses to underlying environmental variation (e.g., habitat filtering). Past assessments have relied on broad metrics such as species richness, which may obscure both pathways. Species richness alone fails to capture well-documented patterns of ecological winners, losers, and functional turnover that occur as native habitats are converted to modified lands (Filgueiras et al. 2021), altering the ecological roles present in a community far more than species counts (Levey et al. 2025).

Functional traits offer a more nuanced representation of biodiversity, capturing how species forage, move, and use resources, and are more directly linked to habitat conditions and ecological processes than richness alone (Bregman et al. 2016; Cadotte et al. 2011). Causally, functional diversity may better capture key mediators of disease risk, such as the composition of predators and competitors, which have been shown to influence how biodiversity directly affects host and vector communities (Sokolow et el. 2015).

Birds are among the most widely used bioindicators in ecology, responding sensitively to habitat quality, vegetation structure, and landscape condition (Anderle et al. 2024; Chiatante et al. 2021; Gregory & van Strien, 2010). Bird functional diversity is especially informative in this context since functionally diverse communities tend to occupy structurally complex, heterogeneous habitats, while functionally homogeneous communities with a narrow range of guilds and trait values are characteristic of disturbed, degraded, or human-modified environments (Levey et al. 2021; Weeks et al. 2025). Since *Ae. aegypti* and *Ae. albopictus* mosquitoes are most prevalent in precisely these disturbed environments due to high occurrence of preferred breeding containers (e.g., anthropogenic water-holding objects; Getis et al. 2003; Shragai & Harrington 2019), bird functional diversity may serve as an informative indicator of the habitat conditions relevant to vector presence, even in the absence of evidence of direct biotic interactions between birds and dengue vectors such as predation (Keesing et al. 2006) and disease reservoir dynamics (Allan et al. 2009).

Among functional diversity metrics, functional dispersion (FDis) measures the mean distance of species from a community centroid in multidimensional trait space, the latter an abundance-weighted measure of how broadly trait variation is distributed across the community that is largely insensitive to species pool size (Laliberté & Legendre 2010; Schleuter et al. 2010). Landscape composition is a key driver of both bird functional diversity (Levey et al. 2025) and *Ae. aegypti* and *Ae. albopictus* mosquito distributions (Kumar et al. 2023; Shragai et al. 2022). Built cover, agricultural land, and forest loss are associated with reduced bird functional diversity (Fahrig, 2003; Newbold et al. 2015) and with increased *Ae. aegypti* and *Ae. albopictus* abundance (Farner et al. 2025; Getis et al. 2003; Simard et al. 2005). If bird FDis integrates information across the habitat conditions relevant to *Aedes* distributions, it could serve as an effective indicator of dengue vector risk. Moreover, if landscape composition is relevant to *Aedes* distributions partly through its effect on bird functional diversity rather than solely through direct habitat modification, FDis may mediate the landscape-mosquito relationship, offering both mechanistic insight and a practical surveillance tool.

Here we test whether bird functional diversity predicts the presence of dengue-transmitting mosquitoes (*Ae. aegypti* and *Ae. albopictus*) across two working landscapes in Costa Rica, and whether the relationship between landscape composition and mosquito presence is mediated through bird functional diversity. We compare six classes of predictor variables, including bird functional diversity indices, bird foraging guild abundance, bird morphometric traits, on-ground vegetation structure, plant diversity, and landscape cover categories derived from remote sensing, in a systematic modelling framework aimed at determining the top predictor variable class and the top predictor variable within the top predictor variable class. We hypothesize that (1) bird functional dispersion will be the top-ranked predictor of *Ae. aegypti* and *Ae. albopictus* adult and larval presence across predictor classes, reflecting shared habitat filtering by landscape condition; (2) landscape composition will be the primary environmental driver of FDis, outperforming fine-scale vegetation metrics; and (3) the effect of landscape on mosquito presence will be mediated through FDis, consistent with a pathway from landscape disturbance to functionally homogeneous bird communities to elevated vector presence. To our knowledge, our study is the first to test whether variation in bird functional diversity predicts the presence probability of dengue vector mosquito species.

## Material and methods

### Study area and site selection

We conducted our study in two Costa Rican landscapes, including in the Quepos canton in Puntarenas province (9°26’18.8“N 84°09’38.3”W) and the Puerto Viejo, Sarapiquí canton in Heredia province (Sarapiquí hereafter; 10°26’45.2“N 84°01’12.9”W) from July to August 2025. The study landscapes are separated by 115 km. To align with our aim to assess the major habitat types of each landscape, we targeted major habitat types within ten sampling polygons that contained ten subsites each, for a total of 100 sampling locations. The habitat composition of the subsites ranged from intensive agricultural land interspersed with rural homes and roads to densely populated urban centers with low native vegetation cover to conserved forested habitat with low urban infrastructure. In both Quepos and Sarapiquí, cattle ranching is a dominant agricultural activity, with plantation agriculture is also present. Differences in agricultural activities between the landscapes include commercial African oil palm (*Elaeis guineensis*) plantations in Quepos and pineapple (*Ananas comosus*) plantations in Sarapiquí. Both *Ae. aegypti* and *Ae. albopictus* occur in Quepos and Sarapiquí, with occurrence reports around residential areas in common household items (tires, plastic bottles, buckets, etc.; Rojas-Araya et al. 2017).

### Larval mosquito sampling

Within each of the 100 sampling locations, we conducted exhaustive searches for water-holding containers that could serve as *Aedes* larval habitat, including any outdoor object capable of retaining water for more than 48 hours. Each water-holding container was visually inspected for the presence of mosquito larvae and pupae. When present, we collected all individuals if fewer than ten were observed. If ten or more were present, we collected a subsample of at least ten individuals. We collected samples using plastic transfer pipettes and preserved them in 50 mL tubes containing approximately 20 mL of 70% ethanol. We identified *Ae*. *aegypti* and *Ae*. *albopictus* larvae using a stereomicroscope and published morphological keys (Chaverri 1995; Rueda 2004). For all containers with *Ae*. *aegypti* or *Ae*. *albopictus* larvae, we recorded container type (e.g., tire, trash receptacle, plant axil).

### Adult mosquito sampling

We collected adult mosquitoes at each sampling location (N=100) using a custom-built backpack aspirator. One trained collector conducted aspirations for 10 min per sampling location, focusing primarily on outdoor microhabitats. At a subset of locations, we conducted indoor aspirations. At the end of each 10 min aspiration, mosquitoes were exposed to ether and transported to the field lab. We identified *Ae*. *aegypti* and *Ae*. *albopictus* adults morphologically using published keys and sorted them by species and sex with a stereomicroscope (Chaverri 1995; Rueda 2004).

### Bird sampling

Within each of the ten sampling polygons, we established three 200-m bird transects on roads separated by at least 200-m (Bibby et al. 2000). We located each bird transect in close proximity (<30 m) to at least 3 mosquito aspiration and larval search locations. Two observers (DRL and JFS) collected all bird data together by recording all birds seen and heard while walking at a slow and steady pace (∼1 km/hr) along the transect route (Bibby et al. 2000). We estimated Euclidean distances to each bird individual to later standardize detection probabilities across habitat types (Thomas et al. 2010).

### Vegetation sampling

We surveyed vegetation and abiotic conditions around each adult and larval mosquito sampling locations after bird sampling using three 5 × 5 m quadrats separated by at least 25 m. We collected plant diversity data and structure data within the sampling space, including: 1) percent cover of ground cover types (bare soil, impervious surfaces, water, grass, leaf litter, shrubby vegetation, and plastic and rubber water-holding containers), 2) a tally of water-holding objects, and 3) percent cover of three canopy layers. We measured percent canopy cover using a fisheye lens when possible; otherwise, estimates were made by eye when low vegetation obstructed the view. We estimated canopy cover at three strata, including a 0–5 m understory and small plant layer, a 6–15 m midstory layer, and a > 16 m upper canopy layer.

### Remote sensing and drone imagery

To quantify landscape variables, we extracted remotely sensed covariates in Google Earth Engine using a standardized 100 m × 100 m square buffer around each sampling location (N=100). We characterized, utilizing 10 m resolution Google Dynamic World V1 and 30 m resolution Landsat 8 collection 2 Level 2 (aggregated across 2020-2025 to improve cloud contamination), the following: 1) built-up area, 2) built-up volume, 3) tree cover, 4) grass cover, 5) crop cover, 6) bare ground cover, 7) pineapple plantation cover, 8) palm oil plantation cover, and 9) surface temperature. For surface temperature measurements, we applied quality filtering using Landsat QA bands and retained only physically plausible surface temperatures; we then summarized temperature within each buffer using the median across all valid observations. Temperature values therefore reflect a long-term relative thermal profile of each site rather than conditions at the specific sampling date.

### Data analysis

We conducted all analyses in R v4.5.3 (R Core Team 2026) and stored data and code on Figshare (https://figshare.com/s/12c7cf4d5598198fa40f; Levey et al. 2026). For analyses, we linked values from each bird transect to the nearby mosquito and vegetation sampling locations. We calculated functional diversity metrics for resident bird communities using the FD package with the *dbFD()* function (Laliberté & Legendre 2010), including functional dispersion (FDis), functional richness (FRic), functional evenness (FEve), functional divergence (FDiv), and Rao’s quadratic entropy (RaoQ). We calculated FDis using morphometric traits, including beak morphology, body size, wing and tail measurements, and foraging and ecological traits, including foraging method, trophic level, trophic niche, and primary lifestyle (AVONET; Tobias et al. 2022). These trait dimensions reflect how birds acquire resources and interact with their environment and are consistently available across species through global trait databases, making them practical for replication. We also gathered breeding traits (Birds of the World 2026) but did not include them in the primary FDis calculation due to stronger evidence of morphometric and foraging traits responding significantly to tropical landscape and habitat gradients (Barros et al. 2019; Levey et al. 2021). We calculated trait distances between species using Gower’s distance to accommodate mixed trait types, and we computed FDis as the abundance-weighted mean distance of species from the community centroid in multidimensional trait space.

To assess whether a simplified trait set could reproduce the signal of the full multi-trait FDis, we calculated FDis separately for the original morphometric and foraging trait classes individually, as well as for breeding traits (breeding months, clutch size, nest height, nest construction, and nesting microhabitat) and a fully expanded trait set combining all three classes (Birds of the World 2026). We fit single-predictor binomial GLMs for each trait-class FDis variant and compared against the original FDis and null models using BIC for each supported response variable.

### Within-class model selection

Within each of six variable classes, including remote sensing landscape metrics, on-ground vegetation and ground structure, plant diversity, bird functional diversity indices, bird morphometric traits, and bird foraging guild abundance, we identified the most informative predictor with a separate univariate binomial generalized linear model (GLM) for each candidate predictor and each response variable. We set response variables as the presence/absence of *Ae. aegypti* larvae, *Ae. albopictus* larvae, *Ae. aegypti* adults, *Ae. albopictus* adults, and combined *Aedes* spp. at larval and adult stages (i.e., either *Aedes* species at either life stage). We standardized all predictors to mean zero and unit standard deviation prior to modelling to allow direct comparison of effect sizes. We compared models using the Bayesian Information Criterion (BIC), implemented in base R with the *BIC()* function, and we retained the predictor with the lowest BIC within each class. Candidate predictors within the remote sensing, vegetation structure from on-ground surveys, and plant diversity classes included both individual variables and their respective composite principal component axes (remote sensing - RS PC1, on-ground survey of vegetation structure - OGS PC1, and plant diversity - PD PC1), allowing the within-class selection procedure to determine whether composite or individual metrics better predicted each response variable. To avoid retaining unstable estimates driven by complete or near-complete separation, we excluded predictors with coefficient estimates or confidence interval bounds exceeding ±10 log-odds prior to within-class selection.

### Cross-class model comparison

We entered the top-ranked predictors from each of the six variable classes, identified through the within-class selection step above, into a cross-class comparison to identify the best overall predictor of mosquito presence. This step compared six single-predictor binomial GLMs, one per class representative, against each other and against a null intercept-only model using BIC, identifying the variable class with the strongest overall support for predicting mosquito presence. Prior to fitting a combined model including all six class representatives simultaneously, we assessed pairwise Pearson correlations among class representative predictors and checked for multicollinearity using variance inflation factors (VIF) implemented in the performance package (Lüdecke et al. 2021). We compared the combined model against the single-predictor models using BIC to assess whether combining class representatives improved on the best single-predictor model. We considered modes preferred over the null when ΔBIC < −2 relative to the null model. We assessed model fit for the best-supported model using the Tjur R² coefficient of discrimination.

### Random effects and spatial structure

We initially specified region and subsite as nested random intercepts in binomial generalized linear mixed models (GLMMs), fit using the lme4 package (Bates et al. 2015), to account for spatial clustering of transects within polygons and polygons within regions. However, models including random effects showed singular variance estimates or were not preferred over simpler GLM specifications by BIC across all response variables, suggesting that bird functional dispersion adequately captured the relevant spatial variation in mosquito presence. The spatial independence assumption was further supported by the minimum separation distances described above. Therefore, we fit final models as binomial GLMs without random effects.

### Mediation analysis

To test whether the effect of landscape composition on mosquito presence operated through bird functional dispersion rather than directly, we conducted a bootstrap mediation analysis. We first confirmed that landscape PC1 predicted FDis (path a) and that FDis predicted mosquito presence after accounting for landscape PC1 (path b) using linear and binomial GLMs respectively. The indirect effect (a × b) was estimated by bootstrapping both path coefficients simultaneously across 1000 resamples of the dataset, and we computed a bootstrapped 95% percentile confidence interval using the boot package (Canty & Ripley 2024). We considered mediation supported when the bootstrapped confidence interval for the indirect effect excluded zero.

## Results

### Biodiversity

We detected a total of 128 bird species (83 in Quepos and 105 in Sarapiquí) and 462 plant species (280 in Quepos and 258 in Sarapiquí) across both study landscapes (Fig. S1). In Quepos, we detected a total of 829 bird and 3902 plant individuals. In Sarapiquí, we detected 615 bird and 5757 plant individuals. The most abundant bird species in Quepos were the Great-tailed Grackle (*Quiscalus mexicanus*), White-winged Dove (*Zenaida asiatica*), Variable Seedeater (*Sporophila corvina*), Tropical Kingbird (*Tyrannus melancholicus*), and Black Vulture (*Coragyps atratus*), while the most abundant bird species in Sarapiquí were the Variable Seedeater, Clay-colored Thrush (*Turdus grayi*), Common Tody Flycatcher (*Todirostrum cinereum*), Great-tailed Grackle, and Tropical Kingbird (Fig. S1). For plants, the species *Elaeis guineensis* (African Oil Palm), *Sphagneticola trilobata*, *Cecropia insignis*, *Guazuma ulmifolia*, and *Phyllanthus niruri* represented the most abundant species, while in Sarapiquí, the species *Pentaclethra macroloba*, *Ananas comosus* (Pineapple), *Colocasia esculenta* (Taro), *Piper auritum*, and *Miconia elata* represented the most abundant species (Fig. S1). For *Ae*. *albopictus* larvae, we detected presence at 43% of urban sampling locations, 29% of rural, and 7% of forest. For *Ae*. *aegypti* larvae, we detected presence at 30% of urban sampling locations and 0% at rural and forest. For *Ae*. *albopictus* adults, we detected presence at 47% of urban sampling locations, 29% of rural, and 21% of forest. For *Ae*. *aegypti* adults, we detected presence at 17% of urban sampling locations, 6% of rural, and 0% of forest.

### Within-class model selection

Within each predictor class, the top-ranked variable differed between *Ae. aegypti* and *Ae. albopictus* for the landscape, vegetation structure, and plant diversity classes, while the top-ranked bird community predictors were consistent across species (Table S1). Within the former three classes, individual variables consistently outperformed their respective composite PC1 axes for all response variables (ΔBIC range: 1.3–5.6), indicating that single land cover or vegetation metrics captured more variation in mosquito presence than multivariate composite indices. For *Ae. aegypti*, the top landscape predictor was built cover, grass cover was the top vegetation structure predictor, and herbaceous plant abundance was the top plant diversity predictor. For *Ae. albopictus*, bare cover, understory cover, and bush plant abundance were the top predictors in those respective classes. Across both mosquito species and life stages, FDis was the top-ranked bird diversity predictor, beak depth was the top bird morphometric predictor, and omnivorous bird abundance was the top bird foraging predictor. Full within-class results for adult stages and combined *Aedes* spp. are presented in Table S1.

### Cross-class model comparison

Pairwise correlations among representatives ranged from r = −0.18 to −0.39, and VIF values for all predictors in the combined model were below 5 across all response variables, indicating acceptably low collinearity. Across all supported response variables, the bird diversity class consistently showed the strongest support relative to the null model, with ΔBIC values ranging from −4.0 to −8.5 (Fig. 2A). Bird foraging diversity was the second most consistently supported class, with negative ΔBIC values across three of the four supported responses. Landscape metrics showed moderate support for *Ae. aegypti* larval and adult presence but not for *Ae. albopictus* larvae and adults. Vegetation structure, plant diversity, and bird morphometric traits showed weak or inconsistent support, with several cells showing positive ΔBIC values. The bird diversity class was the only predictor class preferred over the null for all four supported response variables (Fig. 2A).

**Fig. 1.**
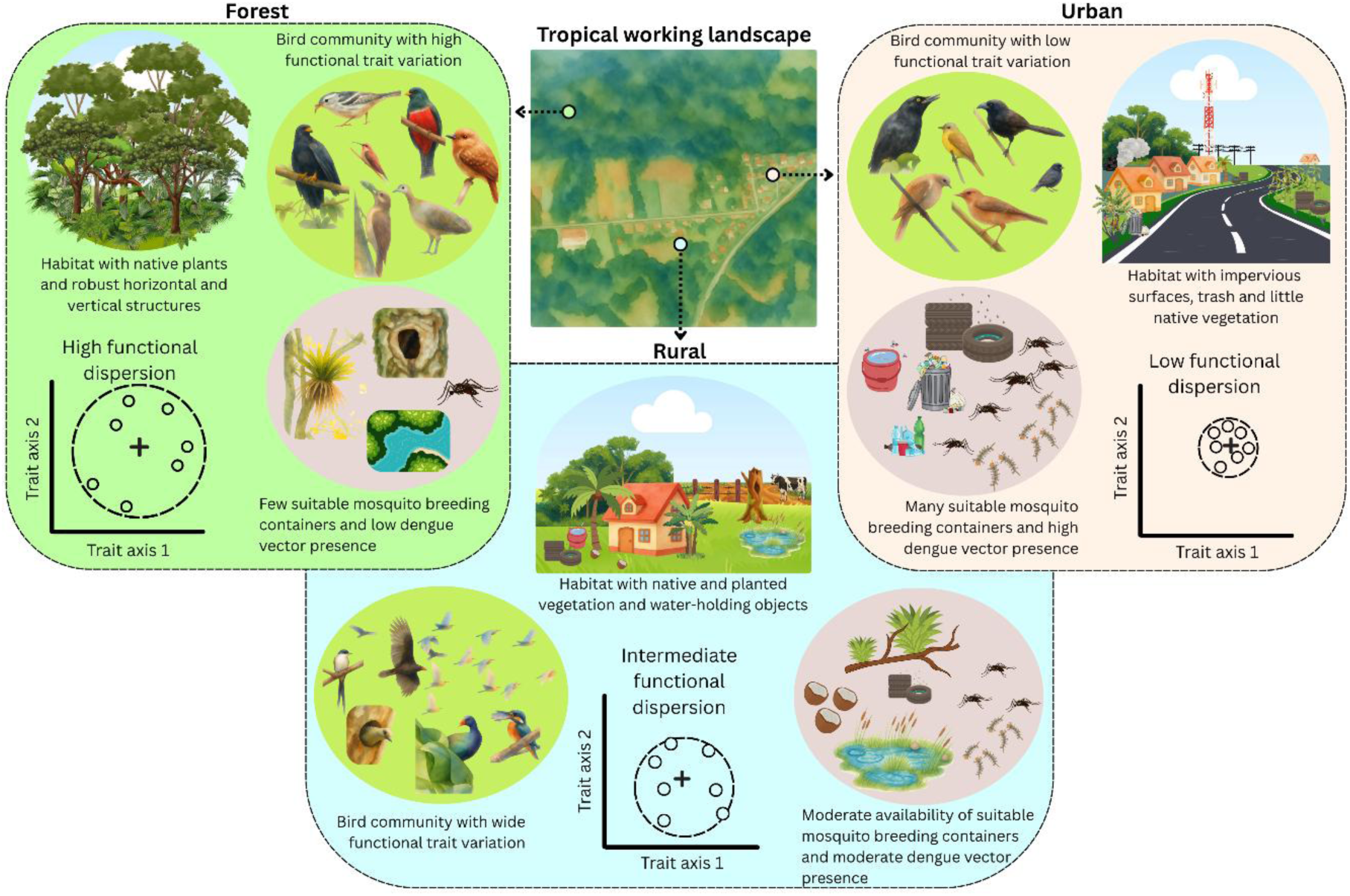
Conceptual illustration of expected relationships between sampling locations within habitat types, bird functional diversity, and dengue vector presence in tropical working landscapes. The central aerial image represents a tropical working landscape encompassing a gradient of habitat types, from forest (left) to rural (center) to urban (right). Within each habitat, bird communities differ in functional trait variation, or the spread of species across morphological, foraging, and breeding trait space, illustrated by trait space schematics showing functional diversity values for sampling sites (points) distributed around a community centroid (+ symbol). Dashed ellipses represent the extent of functional dispersion (FDis): a wide ellipse indicates high FDis (many guilds, varied morphology) and a narrow ellipse indicates low FDis (few guilds, similar morphology). Mosquito breeding habitat availability co-varies with habitat type, with more suitable breeding sites in disturbed environments. Together, high bird functional dispersion and low mosquito breeding site availability are expected to co-occur in structurally complex, native habitats, while low bird functional dispersion and high mosquito breeding site availability are expected in highly modified habitats. Habitat categories, FDis values, and mosquito abundances are illustrative and do not represent observed data from this study.

**Fig. 2.**
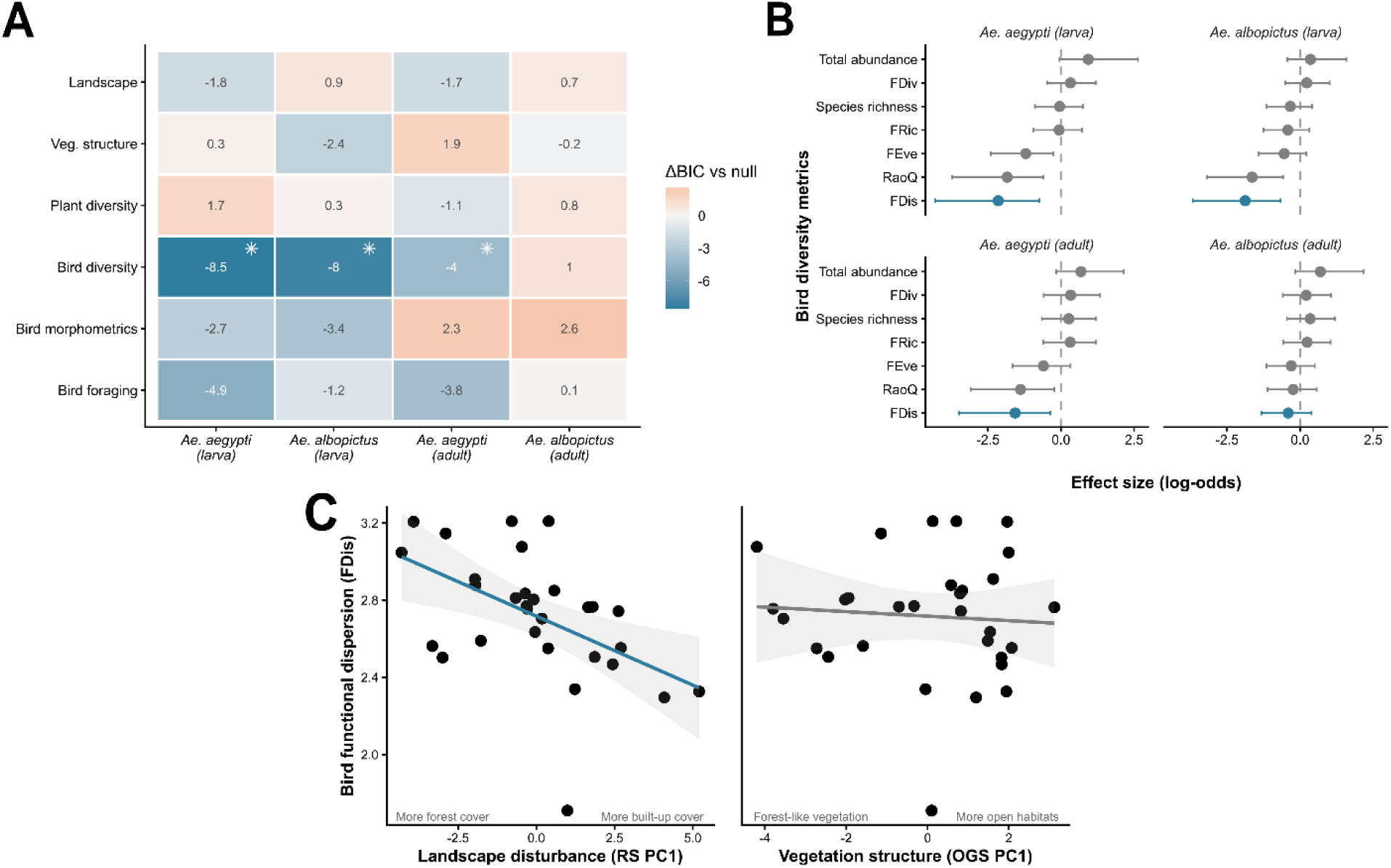
Analytical results linking landscape and vegetation structure and composition, bird functional diversity, and mosquito presence. (**A**) Heatmap showing ΔBIC relative to a null model for the top-ranked predictor within each variable class and each supported response variable. Blue cells indicate models preferred over the null (ΔBIC < 0) and orange cells indicate models not preferred. Asterisks (*) indicate that the bird diversity class was the top-ranked predictor across all classes for that response. (**B**) Effect sizes (log-odds ± 95% CI) of bird diversity indices for each supported response variable. (**C**) Relationship between a landscape-scale gradient PC1 of remote sensing derived variables (left panel - RS PC1) and a habitat-scale vegetation structure gradient PC1 of on-ground survey vegetation variables (right panel - OGS PC1) with bird FDis.

Within the bird diversity class, FDis consistently showed the largest negative effect size and lowest BIC among all bird diversity indices tested (Fig. 2B), outperforming FRic, FEve, FDiv, and RaoQ, and species richness and total abundance across supported responses (Fig. 2B). FDis was also the top-ranked predictor across all six predictor classes for *Ae. aegypti* (BIC = 31.51, ΔBIC = −8.54 relative to null) and *Ae. albopictus* larval presence (BIC = 36.47, ΔBIC = −7.98; Table S1, Fig. 2B), and for adult *Ae*. *aegypti* (BIC = 29.41, ΔBIC = −4.02). The combined model including the top predictor from each class did not improve on the FDis-only model for either larval response, and collinearity values indicated moderate correlation among some class representatives (VIF ≤ 4.34), with FDis showing the lowest collinearity (VIF = 1.47). FDis did not outperform the null model for adult *Ae. albopictus* (ΔBIC = +2.39) or combined adult *Aedes* spp. (ΔBIC = +1.95).

### Trait-class decomposition of FDis

The original FDis, calculated from morphometric and foraging traits, outperformed all trait-class variants for every supported response variable (Table S2). No single trait class reproduced the predictive signal of the original FDis. Morphometric, foraging, and breeding FDis all failed to improve on the null model for most responses. The expanded FDis calculated from all three trait classes combined also performed worse than the original for all supported responses (ΔBIC range relative to original: +0.2 to +4.1). Correlations among FDis variants were moderate (r = 0.48–0.82), with morphometric and foraging FDis showing a weak negative correlation (r = −0.22), suggesting these two trait dimensions capture partially independent axes of functional variation across the study landscapes.

### FDis and mosquito presence

The probability of *Ae. aegypti* larval presence decreased steeply with increasing FDis (β = −2.15, p = 0.015; Fig. 3). A similar negative relationship was observed for *Ae. albopictus* larval presence (β = −1.88, p = 0.011). Combined *Aedes* spp. larval presence showed an identical pattern to *Ae. albopictus* larval presence because *Ae. albopictus* larvae were detected at all sites where any *Aedes* larvae were present (i.e., *Ae. aegypti* larva detections always co-occurred with *Ae. albopictus*, but not vice versa). Consequently, the combined *Aedes* spp. larvae response variable was equivalent to *Ae. albopictus* larval presence. For adult *Ae. aegypti*, the probability of presence also decreased with increasing FDis (β = −1.57, p = 0.043). We detected no significant relationship between FDis and presence for adult *Ae. albopictus* (β = −0.41, p = 0.325) or adult *Aedes* spp. combined (β = −0.47, p = 0.249). The influence of the lowest observed FDis value (FDis = 1.71) was assessed by refitting the *Ae. aegypti* larval model without that observation; the coefficient changed by less than 0.01 log-odds (with: β = −2.149; without: β = −2.139), confirming that this point was not influential (Fig. 3).

**Fig. 3.**
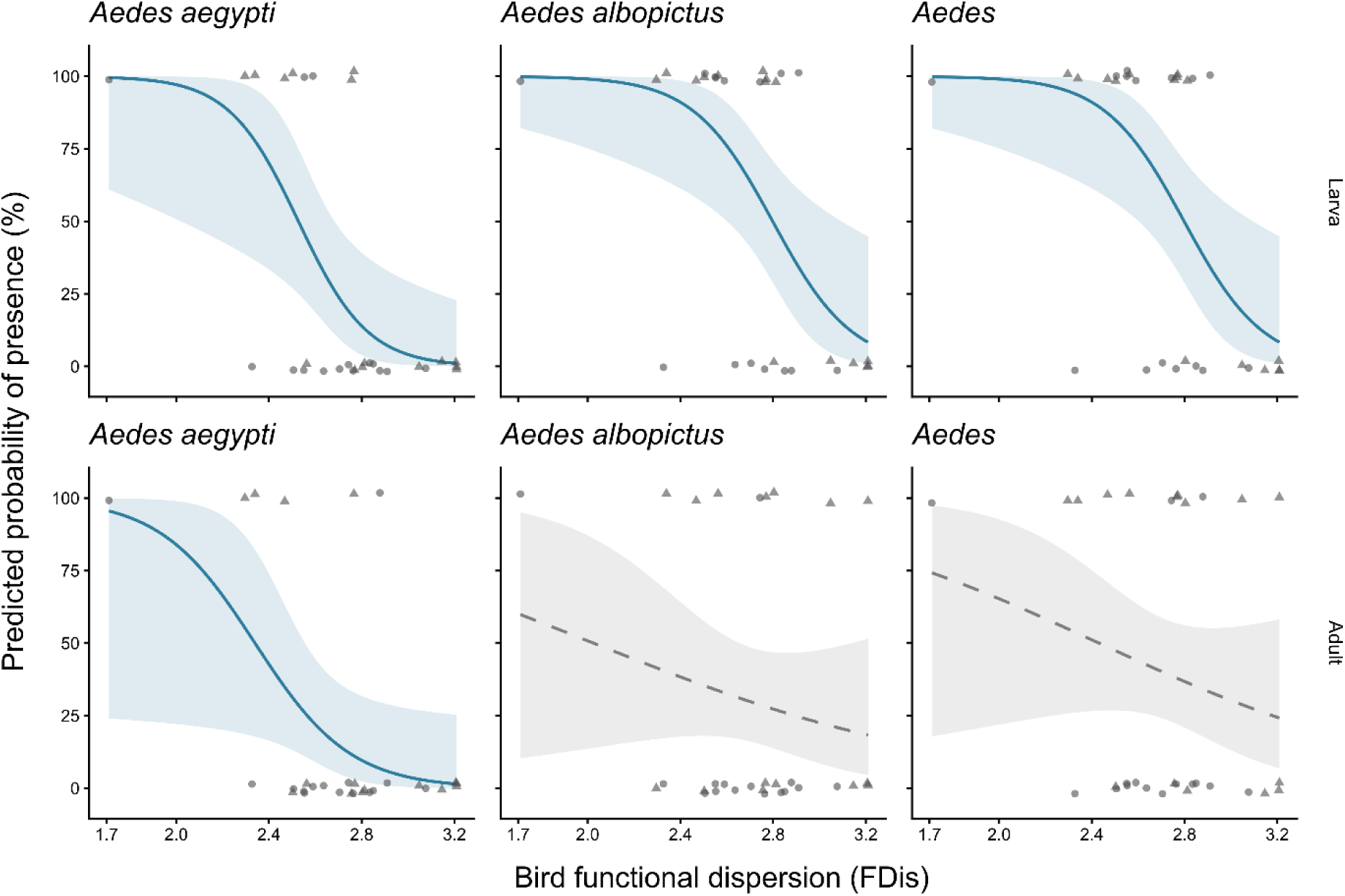
Predicted probability of *Aedes* mosquito presence as a function of bird functional dispersion (FDis) for six response variables across larval (top row) and adult (bottom row) life stages for *Ae. aegypti*, *Ae. albopictus*, and both combined. Blue solid lines and shaded 95% confidence interval ribbons indicate responses for which the FDis model was preferred over the null model (ΔBIC < −2); gray dashed lines and ribbons indicate responses for which FDis was not preferred. Points show observed presence (1) and absence (0) with minor vertical jitter to reduce overlap; point shapes indicate sampling region (circles: Quepos; triangles: Sarapiquí). FDis values range from 1.7 (functionally homogeneous communities) to 3.2 (functionally diverse communities).

### FDis and environmental predictors

Landscape composition (RS PC1) was the top-ranked environmental predictor of bird functional dispersion, outperforming vegetation structure and plant diversity indices by at least 7 BIC units (Fig. 2C, Table S2). Bird functional dispersion decreased with increasing landscape PC1 (β = −0.16, p = 0.004, R² = 0.26), indicating that more disturbed, human-modified landscapes were associated with lower bird functional dispersion. Vegetation structure PC1 and plant diversity PC1 did not improve on the null model for FDis (ΔBIC > 7 relative to the landscape model).

### Mediation analysis

The indirect effect of landscape composition on mosquito presence mediated through bird functional dispersion was supported for *Ae. aegypti* larval presence (bootstrapped 95% CI: 0.39– 8.12) and *Ae. albopictus* larval presence (bootstrapped 95% CI: 0.52–6.48). The indirect effect for adult *Ae. aegypti* presence was positive in direction but associated with high uncertainty (bootstrapped 95% CI: -0.10–109.09). The indirect effect was not supported for *Ae. albopictus* adult presence (bootstrapped 95% CI: −0.50–1.63).

## Discussion

Bird functional dispersion (FDis) was the strongest predictor of *Ae. aegypti* and *Ae. albopictus* presence across two working landscapes in Costa Rica and consistently outperformed landscape composition, vegetation structure, ground cover types, plant diversity, and other bird community metrics in cross-class model comparisons. The probability of larval *Ae*. *aegypti* and *Ae*. *albopictus* presence declined steeply with increasing FDis, and this relationship extended to adult *Ae*. *aegypti*. Landscape composition was the primary driver of FDis, and its effect on mosquito presence was mediated through FDis rather than operating directly. Together these results suggest that standard bird surveys could complement entomological surveillance as a practical, ecologically grounded tool for identifying habitats at elevated dengue vector risk in tropical landscapes.

### Biotic interaction and habitat filtering mechanisms

Bird functional diversity could influence mosquito presence through direct biotic interactions. Insectivorous birds may suppress adult mosquito populations through predation, and functionally diverse communities that exhibit broader foraging strategies may exert more varied predation pressure across microhabitats (Glidden et al. 2021). However, the most parsimonious interpretation is a process of shared habitat filtering by landscape disturbance where both communities respond to the same underlying habitat conditions in opposing directions (Cadotte & Tucker 2017). Human-modified landscapes characterized by high built cover, agricultural land, and reduced native vegetation support functionally homogeneous bird communities dominated by a narrow range of foraging strategies and body sizes (Suárez-Castro et al. 2026), while simultaneously providing the container habitat availability, microclimate conditions, and reduced predator and competitor diversity that favor mosquito disease vectors (Chaves et al. 2021), such as *Aedes* spp. mosquitoes (Dalpadado et al. 2022; Shragai et al. 2019). Functionally diverse bird communities, by contrast, are characteristic of structurally complex habitats with less human presence (Weeks et al. 2025) that are less permissive for *Aedes* spp. establishment and persistence (Farner et al. 2025; Getis et al. 2003; Simard et al. 2005). Since FDis integrates community functionality across multiple trait axes simultaneously, it may capture habitat quality more comprehensively than any single landscape or vegetation metric (Luck et al. 2013), explaining why it outperformed remotely sensed land cover variables that habitat types but not its fine-scale structural complexity. Disentangling these pathways would require experimental manipulation or detailed tracking of bird foraging behavior and mosquito microhabitat use.

### Life stage and species differences

The FDis effect was stronger and more consistent across *Aedes* spp. for larval presence than for adult presence. *Ae*. *aegypti* and *Ae*. *albopictus* larvae are sessile and obligately tied to their aquatic habitat, and their presence at a site reflects local container availability and microhabitat conditions at fine spatial scales (Shragai et al. 2019). Adults of both *Ae*. *aegypti* and *Ae. albopictus* are mobile (Harrington et al. 2005; Maciel-De-Freitas et al. 2007; Moore & Brown 2022), but since *Ae*. *albopictus* may disperse more from natal sites given their affinity for rural habitats with more heterogeneously spaced water-holding containers (Honório et al. 2003), their detected presence may be decoupled from the local conditions that FDis reflects. The strong larval signal suggests FDis tracks fine-scale habitat quality relevant to *Aedes* spp. breeding site availability. From a surveillance perspective, the larval result is particularly useful, as larval presence indicates active local breeding rather than incidental adult movement through a site.

### Landscape mediation

Our results suggest that the landscape-mosquito relationship operates beyond container availability or microclimate differences between land use types, with landscape composition influencing mosquito presence primarily through its structuring effect on bird functional diversity. That FDis may capture this gradient more sensitively than landscape metrics themselves is supported by the distribution of detections across habitat types. We detected *Ae*. *aegypti* larvae exclusively at urban sampling locations, yet FDis predicted presence variation among those urban sites, suggesting sensitivity to fine-scale habitat quality within a broad land use category rather than simply separating urban from non-urban environments. This within-category sensitivity is consistent with the finding that landscape PC1 explained only 26% of the variance in FDis, with the remaining variation likely reflecting fine-scale structural and compositional differences among sites that share a coarse land use classification but differ in vegetation complexity, impervious surface density, or green space characteristics. A similar pattern may apply to *Ae*. *albopictus*, which was detected across urban, rural, and forest sites.

Together these observations suggest that the utility of FDis for surveillance may be greatest precisely where coarse land use classifications are insufficient to discriminate among sites of varying vector risk (Matuoka et al. 2020).

### Bird functional diversity as a biomonitoring tool

Our results have practical implications for dengue vector surveillance. The practical workflow implied by our results is straightforward: conduct point count or transect surveys to characterize the local bird community, calculate FDis with abundance data and morphometric and foraging functional traits, and use the resulting value as an index of relative dengue vector risk across landscapes. Sites with low FDis, which contain functionally homogeneous communities characteristic of disturbed landscapes, would be prioritized for entomological follow-up and vector control. While bird surveys and FDis calculations would not replace entomological surveillance, these techniques may provide a spatially scalable complementary tool for identifying high-risk habitats in data-limited settings. Future studies should test whether the FDis-mosquito relationships generalize across broader geographic contexts, species pools, and seasonal conditions. The consistency of the signal across the two geographically distinct landscapes sampled here suggests it is not idiosyncratic, but broader replication will be needed to establish its generality and develop a predictive model with wider applicability.

Several limitations of the present study warrant acknowledgment. We detected consistent effects across two landscapes despite conducting the study over a single field season. The stability of the FDis-mosquito relationship across seasons and years in Costa Rican landscapes remains to be tested, as dry and wet seasons may influence bird community composition (Hendershot et al. 2020) and availability of mosquito breeding containers (Seidahmed & Eltahir 2016), and climate change is expected to increase disease risk (Childs et al. 2025; Hidalgo et al. 2026). Though we found similar patterns across two distinct landscapes, expanding sampling to additional landscapes, including regions at different elevations and urbanization gradients, will be critical for assessing the generality of these results and for developing a predictive model with broader applicability. Finally, our results establish a correlation between bird functional diversity and mosquito presence but do not demonstrate causation. FDis is best interpreted as a habitat indicator rather than a driver of mosquito suppression. Future studies that explicitly quantify vegetation microstructure, container availability, and bird activity patterns and diet content at the same sites would help disentangle these pathways.

## Acknowledgements

Authors acknowledge Erick López for his technical support during field work and Betzabé Alfaro for support in processing mosquito larvae. This work was funded by the Stanford Doerr School of Sustainability – Disease Ecology in a Changing World Program, the Stanford Sustainability Accelerator, the Stanford King Center on Global Development, and the Stanford Institute for Human-Centered Artificial Intelligence. The work was also supported by University of Costa Rica through project # C5-237, funded by the Vice Rector’s Office for Research. EAM and RLF received additional support from the National Institutes of Health (R35GM133439).

