## Supplementary material for "Bird functional dispersion predicts dengue vector presence across tropical working landscapes in Costa Rica": Table S

**Supplementary materials**

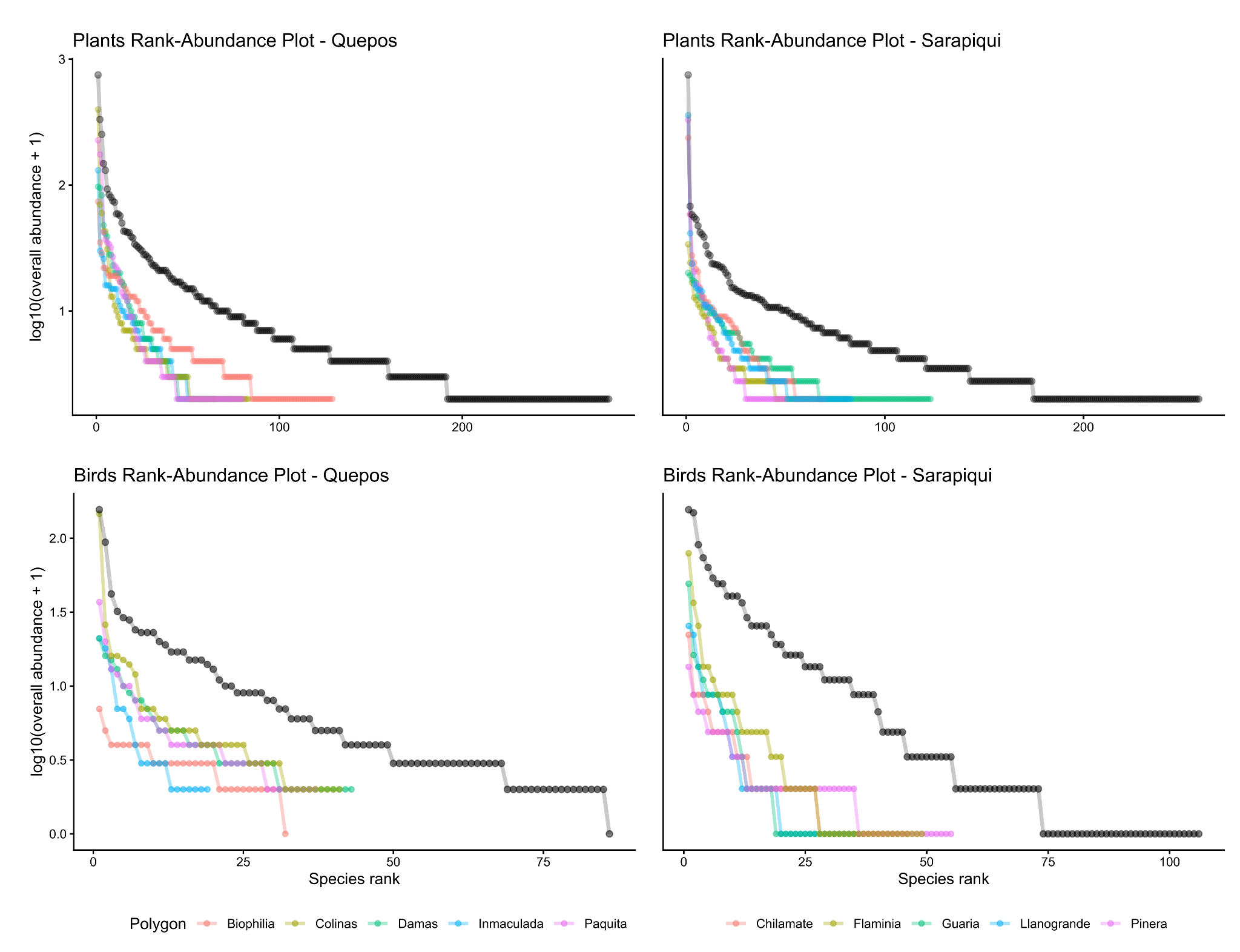

**Fig S1.** Rank-abundance plots of plant and bird communities at each sampling polygon in Quepos and Sarapiquí. Black lines indicate the total combined community across sampling locations. Abundance values are log(x + 1) transformed for improved interpretability. More shallow curves indicate a more even abundance distribution with less dominance of a small number of species, while steeper curves indicate a less even abundance distribution with a small number of dominant species.

**Table S1.** Top-ranked predictor within each variable class for each *Aedes* mosquito response variable, identified through within-class BIC-based model selection. For each variable class, the predictor with the lowest BIC in a univariate binomial GLM was retained as the class representative. Dashes indicate response variables for which no predictor in that class survived the stability filter (coefficient estimates or 95% confidence interval bounds exceeding ±10 log-odds). Bird diversity and bird foraging predictors were consistent across species and life stages, while landscape, vegetation structure, and plant diversity top predictors differed between *Ae. aegypti* and *Ae. albopictus*. Top predictors from each class were carried forward into the cross-class model comparison. FDis: functional dispersion; OGS: on-ground survey.

| **Variable class** | ***Ae. aegypti* larva** | ***Ae. albopictus* larva** | ***Ae. aegypti* adult** | ***Ae. albopictus* adult** | ***Aedes* spp. larva** | ***Aedes* spp. adult** |
| --- | --- | --- | --- | --- | --- | --- |
| **Remote sensing** | Built cover | Bare cover | Built cover | — | Bare cover | — |
| **Vegetation structure** | Grass cover | Understory cover | — | — | Understory cover | — |
| **Plant diversity** | Herbaceous plant abundance | Bush plant abundance | Bush plant abundance | Herbaceous plant abundance | Bush plant abundance | Herbaceous plant richness |
| **Bird diversity** | FDis | FDis | FDis | — | FDis | FDis |
| **Bird traits** | Beak depth | Beak depth | Secondary feather length | Nares length | Beak depth | Hand-wing index |
| **Bird foraging** | Omnivore abundance | Omnivore abundance | Omnivore abundance | Omnivore richness | Omnivore abundance | Omnivore richness |

**Table S2.** Comparison of binomial GLMs predicting *Aedes* spp. Mosquito presence using bird functional dispersion (FDis) calculated from different trait class combinations. For each response variable, FDis was calculated separately from morphometric traits (beak morphology, body size, wing and tail measurements), foraging and ecological traits (foraging method, trophic level, trophic niche, and primary lifestyle), breeding traits (breeding months, clutch size, nest height, nest construction, and nesting microhabitat), and a complete trait set combining all three classes. The original FDis used in primary analyses contained morphometric and foraging traits only, based on previous studies demonstrating strong responses of bird morphometrics and foraging trait diversity to tropical land use change. We compared models with Bayesian Information Criterion (BIC); we calculated ΔBIC values relative to the null intercept-only model and relative to the original FDis model. Lower BIC values indicate better supported models, and we considered models with ΔBIC < -2 relative to the null preferred. The original FDis outperformed all trait-class variants for every supported response variable, suggesting that predictive power derives from integrating multiple trait dimensions rather than any single functional axis.

| **Response** | **Model** | **BIC** | **BIC vs null** | **BIC vs original** | **Best model** |
| --- | --- | --- | --- | --- | --- |
| *Ae. aegypti* (larva) | Null | 40.05 | 0 | 8.54 |  |
| *Ae. aegypti* (larva) | FDis (original - morphometric + foraging) | 31.51 | -8.54 | 0 | Yes |
| *Ae. aegypti* (larva) | FDis (all traits) | 33.73 | -6.32 | 2.22 |  |
| *Ae. aegypti* (larva) | FDis (morphometric only) | 41.6 | 1.55 | 10.09 |  |
| *Ae. aegypti* (larva) | FDis (foraging only) | 37.62 | -2.43 | 6.11 |  |
| *Ae. aegypti* (larva) | FDis (breeding only) | 38.49 | -1.56 | 6.98 |  |
| *Ae. albopictus* (larva) | Null | 44.46 | 0 | 7.99 |  |
| *Ae. albopictus* (larva) | FDis (original - morphometric + foraging) | 36.47 | -7.99 | 0 | Yes |
| *Ae. albopictus* (larva) | FDis (all traits) | 37.31 | -7.15 | 0.84 |  |
| *Ae. albopictus* (larva) | FDis (morphometric only) | 42.62 | -1.84 | 6.15 |  |
| *Ae. albopictus* (larva) | FDis (foraging only) | 45.67 | 1.21 | 9.2 |  |
| *Ae. albopictus* (larva) | FDis (breeding only) | 44.37 | -0.09 | 7.9 |  |
| *Ae. aegypti* (adult) | Null | 33.43 | 0 | 4.02 |  |
| *Ae. aegypti* (adult) | FDis (original - morphometric + foraging) | 29.41 | -4.02 | 0 | Yes |
| *Ae. aegypti* (adult) | FDis (all traits) | 33.57 | 0.14 | 4.16 |  |
| *Ae. aegypti* (adult) | FDis (morphometric only) | 34.57 | 1.14 | 5.16 |  |
| *Ae. aegypti* (adult) | FDis (foraging only) | 35.33 | 1.9 | 5.92 |  |
| *Ae. aegypti* (adult) | FDis (breeding only) | 34.44 | 1.01 | 5.03 |  |
| *Ae. albopictus* (adult) | Null | 40.05 | 0 | -2.39 | Yes |
| *Ae. albopictus* (adult) | FDis (original - morphometric + foraging) | 42.44 | 2.39 | 0 |  |
| *Ae. albopictus* (adult) | FDis (all traits) | 43.03 | 2.98 | 0.59 |  |
| *Ae. albopictus* (adult) | FDis (morphometric only) | 43.39 | 3.34 | 0.95 |  |
| *Ae. albopictus* (adult) | FDis (foraging only) | 42.91 | 2.86 | 0.47 |  |
| *Ae. albopictus* (adult) | FDis (breeding only) | 41 | 0.95 | -1.44 |  |
